# Speed breeding in growth chambers and glasshouses for crop breeding and model plant research

**DOI:** 10.1101/369512

**Authors:** Sreya Ghosh, Amy Watson, Oscar E. Gonzalez-Navarro, Ricardo H. Ramirez-Gonzalez, Luis Yanes, Marcela Mendoza-Suárez, James Simmonds, Rachel Wells, Tracey Rayner, Phon Green, Amber Hafeez, Sadiye Hayta, Rachel E. Melton, Andrew Steed, Abhimanyu Sarkar, Jeremy Carter, Lionel Perkins, John Lord, Mark Tester, Anne Osbourn, Matthew J. Moscou, Paul Nicholson, Wendy Harwood, Cathie Martin, Claire Domoney, Cristobal Uauy, Brittany Hazard, Brande B. H. Wulff, Lee T. Hickey

## Abstract

To meet the challenge of feeding a growing population, breeders and scientists are continuously looking for ways to increase genetic gain in crop breeding. One way this can be achieved is through “speed breeding” (SB), which shortens the breeding cycle and accelerates research studies through rapid generation advancement. The SB method can be carried out in a number of ways, one of which involves extending the duration of a plant’s daily exposure to light (photoperiod) combined with early seed harvest in order to cycle quickly from seed to seed, thereby reducing the generation times for some long-day (LD) or day-neutral crops. Here we present glasshouse and growth chamber-based SB protocols with supporting data from experimentation with several crop species. These protocols describe the growing conditions, including soil media composition, lighting, temperature and spacing, which promote rapid growth of spring and winter bread wheat, durum wheat, barley, oat, various members of the *Brassica* family, chickpea, pea, grasspea, quinoa and the model grass *Brachypodium distachyon*. Points of flexibility within the protocols are highlighted, including how plant density can be increased to efficiently scale-up plant numbers for single seed descent (SSD) purposes. Conversely, instructions on how to perform SB on a small-scale by creating a benchtop SB growth cabinet that enables optimization of parameters at a low cost are provided. We also outline the procedure for harvesting and germinating premature wheat, barley and pea seed to reduce generation time. Finally, we provide troubleshooting suggestions to avoid potential pitfalls.

## 2. Introduction

To improve the productivity and stability of crops there is pressure to fast-track research and increase the rate of variety development. The generation time of most plant species represents a bottleneck in applied research programs and breeding, creating the need for technologies that accelerate plant development and generation turnover. Recently we reported protocols for ‘speed breeding’ (SB), which involve extending the photoperiod using supplementary lighting and temperature control, enabling rapid generation advancement in glasshouses with sodium vapour lamps (SVL) or growth chambers fitted with a mixture of metal halide and light-emitting diode (LED) lighting^1^. By adopting a 22-hour photoperiod and controlled temperature regime, generation times were significantly reduced for spring bread wheat (*Triticum aestivum*), durum wheat (*T. durum*), barley (*Hordeum vulgare*), chickpea (*Cicer arietinum*), pea (*Pisum sativum*), canola (*Brassica napus*), the model grass, *Brachypodium distachyon* and the model legume, *Medicago truncatula*, in comparison to the field or a glasshouse with no supplementary light. Under the rapid growth conditions, plant development was normal, plants could be easily crossed (wheat and barley), and seed germination rates were high. We also demonstrated that SB can be used to accelerate gene transformation pipelines and adult plant phenotyping could be performed under SB conditions for traits such as flowering time, plant height, and disease resistance in wheat, leaf sheath glaucousness in barley, and pod shattering in canola^1^.

The use of extended photoperiod to hasten plant growth is not novel. Sysoeva et al. (2010)^2^ provides an extensive review of the literature surrounding this subject, published within the last 90 years, which outlines successful attempts using spring wheat, barley, pea, chickpea, radish (*Raphanus sativus*), alfalfa (*Medicago sativa*), canola, flax (*Linum usitatissimum*), arabidopsis (*Arabidopsis thaliana*), apple (*Malus domestica*) and rose (*Rosa × hybrida*), among others. More recent examples of photoperiod manipulation to hasten flowering time of crop species include lentil (*Lens culinaris*)^3,4^, pea (*P. sativum*), chickpea (*C. arietinum*), faba bean (*Vicia faba*), lupin (*Lupinus angustifolius*)^5^ and clover (*Trifolium subterraneum*)^6^.

Here, we provide a standardised SB protocol for use in a glasshouse, or a growth chamber with additional data-supported modifications. We provide details for scaling-up plant numbers in the glasshouse, suitable for single seed descent (SSD) to generate large populations. Since plant species, indeed even cultivars within a species, are highly diverse in their response to photoperiod, a universal protocol for all plant species and traits is not possible. We therefore provide instructions for building a low-cost benchtop SB cabinet with controlled lighting and humidity monitoring, suitable for small-scale research projects and trialling SB parameters. Notwithstanding, we have observed that the protocols are flexible and can be tailored to fit a wide range of breeding or research objectives and crop species. By sharing these protocols, we aim to provide a pathway for accelerating crop research and breeding challenges.

### Overview of the procedure

In this protocol, we describe how to implement SB in temperature-controlled glasshouses using supplementary LED lighting, which provides significant cost savings over traditional SVLs. The protocols have been tested in the UK and Australia, with lights from the same company, but with slightly different models. We also outline compatible soil mixes for various crops when growing them under these lighting regimes, along with advice for early harvest to reduce generation time further. We provide supporting data to demonstrate the suitability of these setups to significantly decrease the number of days to flowering and overall generation advancement for spring wheat, barley, canola, chickpea, pea, *B. distachyon*, *M. truncatula,* oat (*Avena strigosa*), grasspea (*Lathyrus sativus*) and quinoa (*Chenopodium quinoa*). We also include the design, step-by-step construction protocol, and operation of a small growth cabinet, which allows control over the light quality, intensity and photoperiod to help optimize the SB recipe for different crops and cultivars before implementing a large-scale glasshouse experiment.

Crop breeding programs commonly use SSD for several generations, on large numbers of segregating plants, to generate homozygous lines with fixed traits^7^. A glasshouse is often preferred for SSD because plant populations can be grown year-round. This process involves both a large investment in time as well as space within the glasshouse. Following the crossing of two homozygous lines, six generations of self-pollination are required to produce progeny that are 98.4% homozygous, which, at a rate of two generations per year, would take three years to complete. While only one or two seeds are needed from each plant to begin the next generation, plant researchers and breeders seek to maximise the number of plants within a restricted space. Plant density can be scaled-up under SB to enable concurrent rapid cycling of large plant populations, which is ideal for SSD programs. To demonstrate this, we evaluated spring wheat and barley sown at different plant densities in a glasshouse fitted with LED supplementary lighting. By comparing the physiological, morphological and yield parameters, we illustrate the normal development of these plants and highlight how this SB approach can save time and resources for SSD programs.

### Development of the protocols

The SB concept was inspired by the efforts of NASA to grow crops in space, using an enclosed chamber and extended photoperiod^8^. In recognising the opportunity to more rapidly produce adult wheat and barley plants and allow faster selection and population development, SB became the norm in cereal research activities at the University of Queensland (UQ), Australia, thanks to Dr Ian Delacy and Dr Mark Dieters. The original protocol was first described and implemented for wheat^9^ and peanut (*Arachis hypogaea*)^10^. Variations of this protocol have been demonstrated to be an efficient system for rapid screening of wheat germplasm for adult plant resistance to various diseases^11-14^ and also for pyramiding multiple disease resistance in barley^15^. The protocol has also been adapted for high-density plant production systems for SSD programs. The current SB protocol described in this paper was developed from the initial implementation described for wheat to include a two-hour dark period that improved plant health^1^. This change was made following experiments in a controlled environment chamber at the John Innes Centre (JIC), UK, and was demonstrated to be suitable for accelerating research activities involving adult plant phenotyping, genetic structuring, and molecular studies like gene transformation in wheat and barley. It was further demonstrated to be suitable for rapid generation advancement for durum wheat (*T. durum*), pea, the model grass, *B. distachyon* and the model legume, *M. truncatula*, and could be scaled up in the SB glasshouse system at UQ, to be made suitable for rapid generation advancement of wheat, barley, canola and chickpea.

### Comparison with other approaches

Perhaps the most well-known strategy to increase generation turnover is ‘shuttle breeding’, introduced by Dr Norman Borlaug in the 1950s at the international Centre for Maize and Wheat Improvement (CIMMYT), which enabled growing two generations per year by sowing wheat populations at field locations differing in altitude, latitude, and climate in Mexico^16^. There is also a long history of extensive efforts to accelerate plant growth of many species by manipulating photoperiod under artificial conditions, as briefly outlined above.

Supplementary lighting is not the only basis for rapid generation advance in plants. A common approach involves exerting physiological stress to trigger flowering and earlier setting of seed. This involves restricting plant growth area (by growing plants at high densities) or nutrient and water access^17^, accompanied by thinning of the plant canopy. Such a method is well-established and documented for rice^18^ and has also been demonstrated for pea (Supplementary Figure 1). Embryo rescue is another common feature in many rapid cycling methods where immature seed is harvested and induced to germinate on culture media, with or without the addition of plant growth regulators (PGR), to negate the waiting time for seed to mature. Bermejo et al. (2016)^19^ used PGR in embryo culture media to promote germination of immature lentil seed to achieve 4 generations annually. Mobini et al. (2015)^20^ sprayed lentil and faba bean plants with PGR to promote early flowering and applied embryo rescue with PGR-enriched agar media to achieve up to 8 and 6.8 generations per year, respectively. Application of PGR is not required for SB, which may be desirable considering the additional time and effort required for handling these and working out the logistics of their application at specific times. In addition, if a species-specific protocol is not available, extensive testing would be needed to optimise such applications. There are also examples of embryo rescue without PGR to shorten generation time. Zheng et al. (2013)^21^ and Yao et al. (2017)^22^ reported up to 8 generations per year for wheat and Zheng et al. (2013)^21^ reported up to 9 generations per year for barley. Both Ochatt et al. (2002)^23^ and Mobini and Warkentin (2016)^5^ reported up to 6.9 and 5.3 generations of pea per year respectively, and Roumet and Morin (1997)^24^ reported 5 cycles per year in soybean (*Glycine max* L.), all with embryo rescue without PGRs. On the other hand, SB conditions without embryo rescue is capable of producing 6 generations per year for spring wheat, barley, chickpea and pea, and 4 generations per year for canola^1^. Testing is needed for any plant species prior to implementation, but this approach is promising for other cereal, pulse and legume crops. Seed of wheat and barley produced under SB conditions can be harvested prematurely at two weeks post-anthesis, followed by a short period of drying and chilling to achieve high and uniform germination rates and healthy plants^1^. Protocols involving embryo rescue are important and useful for breeding and research programs if the required infrastructure is available^25^, particularly for species that are recalcitrant to other parameters used to accelerate generation advancement such as temperature or photoperiod manipulation^26-28^. In comparison, the SB protocols outlined here are less labour intensive, especially with large populations, and laboratory facilities are not required, making the protocols more accessible.

Plant growth can also be promoted by increasing the CO_2_ concentration. For example, for C_3_ plants like rice and wheat, photosynthetic efficiency increases with increasing CO_2_ levels, leading to an increase in biomass and early flowering. In fact, there are documented methods for rapid generation advance in rice that combine restricted root growth and canopy thinning with high CO_2_ concentration, followed by early harvest and embryo rescue to cut down generation times of many rice varieties^29^.

Doubled haploid (DH) technology, where haploid (*n*) embryos are rescued and undergo chromosome doubling (2*n*), is extensively and routinely used in the breeding of several crop species, thus reducing the number of generations required to achieve homozygous lines from six or more to just two generations^30^. Despite this, DH technology has some disadvantages: it can be expensive, requires specialist skills, restricts recombination to a single round of meiosis, and has a variable success rate that may be genotype-dependant^31^. The approach can also be labour intensive for large populations, especially those requiring removal of the embryos from the seed coat. Notably, there is the potential for SB to further accelerate the production of DH lines by speeding up the crossing, plant regeneration and seed multiplication steps.

We have presented a design for building a low-cost benchtop growth cabinet to trial SB. Compared to other published protocols for self-made growth chambers^32,33^, our design makes use of a more widely available control system using a Raspberry Pi and compatible sensors, with codes for the user interface (UI) freely available from GitHub (https://github.com/PhenoTIPI/SpeedSeed3/wiki). The cabinet was trialled for the 22-hour SB lighting, temperature and photoperiod regime (22 °C/17 °C (22 hours/2 hours)), and successfully reproduced the accelerated development of one rapid-cycling variety of each of wheat and pea (Supplementary Tables 1, 2). The component costs for constructing such a cabinet are provided in Supplementary Table 3).

### Limitations of the approach

Different plant species can have markedly different responses when exposed to extended photoperiods. For long-day (LD) plants, time to flowering is often accelerated under extended photoperiods since the critical day length is generally exceeded. This is also the case with day-neutral plants, where flowering will occur regardless of the photoperiod. In contrast, short-day (SD) plants require the photoperiod to be below the critical daylength to flower^34^, which could be at odds with SB conditions. However, there are exceptions and some species show a facultative response where, although flowering is promoted by a particular photoperiod, flowering will still occur in the opposite photoperiod. Furthermore, the time difference between being a SD or LD plant can be a matter of minutes^35^. These factors highlight both a limitation of SB and a point of flexibility. In cases where the photoperiod response is unknown or complex in nature, experimentation of light and temperature parameters is required to optimise a SB strategy, for example, by using the benchtop growth cabinet. For instance, applying extended light prior to and following a shortened photoperiod to induce flowering, could hasten initial vegetative growth and accelerate maturity, respectively, thus producing an overall shorter generation time. Such an approach has been successfully applied to amaranth (*Amaranthus* spp. L), a SD species, where a 16-hour LD photoperiod was used to initiate strong vegetative growth after which plants were transferred to an 8-hour SD photoperiod to induce flowering^36^. The overall effect was a shorter lifecycle and ability to produce eight generations per year rather than two in the field. The need for vernalisation, such as in winter wheat, creates a situation similar to above. Young plants require chilling for a number of weeks to trigger the transition to flowering. Once the vernalisation requirement is met in winter wheat, exposing the plants to extended photoperiod is likely to accelerate growth^37,38^. Overall, the ‘SB recipe’ is more straight forward and easier to implement for LD and day neutral species which do not require vernalisation. Experimentation and optimisation of parameters are highly recommended for each species.

The SB protocols presented here take place in an enclosed, artificial environment, which differs significantly from the field where eventual crop production may occur. While this is acceptable for many activities, such as crossing, SSD and screening for some simple traits^1^, other activities, such as selection for adaptation in the target environment must still occur in the field. Nevertheless, programs alternating between SB and the field save time overall. The ability to shorten generation time further through early harvest of immature seed can interfere with the phenotyping of some seed traits. For this reason, in spring wheat breeding programs where dormant and non-dormant genotypes need differentiating, phenotyping grain dormancy under SB conditions is limited to only four generations per year^9^.

The initial investment to build a glasshouse or purchase a growth chamber with appropriate supplementary lighting and temperature control capabilities is substantial if these facilities are not already available. However, depending on the budget of the research or breeding program, the benefits may outweigh the costs. For instance, an economic analysis performed by Collard et al. (2017)^39^ compared the rapid generation advance (i.e., no phenotypic selection at each generation) with the pedigree-based breeding method (i.e., with phenotypic selection at each generation) for rice and determined that rapid generation (achieved through restricted soil access and canopy thinning) was more cost-effective and advantages would be realized after one year even if new facilities were constructed. Nevertheless, most breeding programs have pre-existing glasshouse facilities that can be converted for SB applications, but careful selection of energy efficient lighting and temperature control systems are needed to minimise operating costs. Research activities often do not require the high plant numbers needed in breeding, so growth chambers are common. The cost of these start at tens of thousands of dollars, making them inaccessible for many projects and a barrier for implementing SB. In addition, the energy to provide extended supplementary lighting is significant. A cost-benefit analysis should be carried out to determine feasibility although there are areas where cost-savings can be made. Supplemental LED lighting provides more efficient power usage and reduced heat than other lighting types, such as SVLs. An estimate of the maintenance and energy costs associated with LED lighting is provided in the supplementary material of Watson and Ghosh et al. (2018)^1^. Investing in solar panels is another strategy to offset the increased energy costs, depending on availability and location.

The investment in SB needs to be weighed in terms of the potential benefits to variety development and research output. As with most technologies, determining the optimal way to integrate SB in a crop improvement program needs careful consideration and may require significant re-design or restructure to the overall program. Prior to implementing such changes, computer simulations are a good way to evaluate the different breeding programs incorporating SB.

### Experimental Design

To set-up an effective SB system, certain factors require careful consideration. These include:

#### a) Lighting requirements

Many lighting sources are appropriate for SB, including SVLs and LEDs^1^. Even incandescent lighting has been shown to accelerate flowering in clover^6^. However, selection should be based on the space available, plant species and energy resources. For example, LED lighting may be preferred due to its energy efficiency although simple incandescent lighting may be suitable within a smaller area, with sufficient cooling to counteract the higher heat output. Plant species may also differ in their response to the different spectra of wavelengths emitted by different lighting sources so this should be carefully considered. The lighting setup for glasshouses and growth chambers detailed in this protocol can act as a starting point but is by no means the final conditions that may be optimum for another situation. The protocols outlined here have been successful for the species trialled but a modified approach may be more suitable for another crop. We recommend mining existing literature and studies on suitable light spectra (particularly with regard to blue to red ratios, red to far-red ratios, and the proportional level of UV light that may be introduced into the system) for the crop and trait of interest.

#### b) Initial light calibrations

Requirements in terms of light quality and intensity for a particular species, cultivar of that species, and desired phenotype, should be determined prior to application on a large scale or use within an experiment. Several ‘dummy’ or ‘test’ growth cycles are recommended to initially assess the rate of growth and quality of the plants so that alterations can be made to enable optimal outcomes. For this purpose, we recommend starting with the benchtop growth cabinet option – the costs of which are low enough to build several and trial, in parallel, different light-combinations, photoperiods and temperatures to determine the optimal conditions to implement on a larger scale, such as a glasshouse, for your crop and trait.

#### c) Germplasm

As detailed above, not all plant species (or indeed cultivars within a species) are amenable to extended photoperiod. Care should therefore be exercised in selection of the germplasm to be grown under SB and appropriate modifications implemented to ensure optimal conditions for each species.

#### d) End-use requirements

The intended end-use of the resultant plants can affect all aspects of the initial set-up of the SB protocol, such as glasshouse space and sowing density. For example, within an SSD program large numbers of plants are grown within a defined space, so an appropriate sowing density needs to be determined. Conversely, a small number of plants needed for a research experiment under variable lighting parameters is more appropriate for a small growth chamber experiment with flexible settings.

#### e) Control conditions

Before beginning a SB experiment, it is important to have replicates of your germplasm growing under the conditions you would normally use in your breeding program or institute. This will allow you to directly compare plant growth parameters (including generation time), operational costs (e.g. electricity) and plant quality. For popular varieties grown for many generations in the field or glasshouses, the control data may be readily available.

## 3. Materials

### Reagents

#### a) Soil

Soil mixtures which have previously been shown to work for certain crops in SB conditions are provided in <u>Table 1</u>. Details of the soil mixture composition can be found in Supplementary Tables 4, 5 and 6.

**Table 1.** List of soil mixes that have been demonstrated to be compatible for speed breeding using our protocols.

| Species | Compatible soil mixes |
| --- | --- |
| Bread wheat (*T. aestivum*) | JIC Cereal Compost Mix, UQ Compost Mix |
| Durum wheat (*T. durum*) | JIC Cereal Compost Mix, UQ Compost Mix |
| Barley (*H. vulgare*) | JIC Cereal Compost Mix, UQ Compost Mix |
| Pea (*P. sativum*) | JIC Cereal Compost Mix |
| Chickpea (*C. arietinum*) | UQ Compost Mix |
| *Brassica rapa* | JIC Cereal Compost Mix |
| *Brassica oleracea* | JIC Cereal Compost Mix |
| Canola (*Brassica napus*) | JIC Cereal Compost Mix, UQ Compost Mix |
| Quinoa (*C. quinoa*) | JIC Peat and Sand Mix |
| Oat (*A. strigosa*) | JIC Cereal Compost Mix |
| Grasspea (*L. sativus*) | JIC Cereal Compost Mix |
| *Brachypodium distachyon* | JIC Cereal Compost Mix, 50% JIC Cereal Compost Mix + 50% JIC Peat and Sand Mix |
| Medicago | JIC Cereal Compost Mix |

#### b) Nutrient feed

Depending on the size of the pots and the type of soil, the plants may need a nutrient feed. If the pots are small (~100 ml), a single or fortnightly application of a liquid nutrient feed should be considered to prevent the plant leaves from turning yellow prematurely with concomitant reduced vigour and seed set. In the JIC glasshouses and growth chambers, we have successfully used Solufeed 1-1-1 from Vitax (http://www.vitaxgrower.co.uk/product/vitafeeds/) for wheat growing in high density trays.

##### Critical

Due to the rapid growth of plants under SB, fertiliser application and swift amelioration of nutrient deficiencies are of utmost importance. Appropriate slow-release fertiliser within the soil media is recommended for growth to maturity, and maintenance of soil pH is important to avoid restriction of nutrient absorption; e.g. a pH that is too acidic can inhibit calcium uptake. Foliar fertiliser applications may be required for rapid access of nutrients to the leaves although some level of calcium deficiency is common. See Supplementary Figure 2 for common symptoms of calcium deficiency. In our experience, for wheat, barley and *Brachypodium*, symptoms are more common at early growth stages during the period of prolific vegetative growth and are relieved at later growth stages. See Troubleshooting (Section 6) for specific suggestions on calcium applications.

### Equipment

The sections below describe the equipment needed for different SB purposes:

Section a: Provides information to set up SB in an existing plant growth chamber or controlled environment room (CER). This section outlines the core “recipe” for programing an existing growth room to set up SB conditions.
Section b: Provides details for the design and construction of a small benchtop cabinet for SB, which may be used for small-scale pilot trials before investing in a larger system, such as a glasshouse. The cabinet has a footprint of 0.225 m^2^ and comfortably accomodates eight 1 L square pots.
Section c: Provides details for setting up SB in a glasshouse using LED lamps for supplementary lighting. Its efficacy is demonstrated for a range of crop species, along with some examples of how single-seed descent for wheat and barley can be carried out. The LED supplemental lighting within glasshouses at JIC (UK) and UQ (Australia), were supplied by the same company, Heliospectra (Göteborg, Sweden). Details of both setups are provided, along with the results of experiments carried out at both locations.

#### Section a) Speed breeding setup

##### i) Lights

We have shown in our previous studies^1^, that any light that produces a spectrum which reasonably covers the photosynthetically active radiation (PAR) region (400-700 nm), with particular focus on the blue, red and far-red ranges, is suitable to use for SB. The referenced study has several examples of these spectra, and similar examples of possible SB spectra are provided here. An appropriate spectral range can be achieved through LEDs, or a combination of LEDs and other lighting sources (e.g. halogen lamps), or in the case of a glasshouse, by simply supplementing the ambient lighting with LEDs or SVLs. We highly recommend that measurements of the light spectrum are taken prior to commencement of the SB experiment. In addition to controlling the light quality, we recommend a photosynthetic photon flux density (PPFD) of approximately 450-500 μmol·m^-2.^s^-1^ at plant canopy height. Slightly lower or higher PPFD levels are also suitable. Crops species vary in their response to high irradiance. However, the suggested level of 450-500 μmol·m^-2.^s^-1^ has been demonstrated to be effective for a range of crop species^1^.

##### ii) Photoperiod

We recommend a photoperiod of 22 hours with 2 hours of darkness in a 24-hour diurnal cycle. Continuous light is another option, but our experience has shown that the dark period slightly improves plant health. Gradually increasing light intensity to mimic dawn and dusk states should be done, if possible, but is not vital. In our previous paper, we have also provided an example where an 18-hour photoperiod was sufficient to achieve faster generation times for wheat, barley, oat and triticale^1^.

##### iii) Temperature

The optimal temperature regime (maximum and minimum temperatures) should be applied for each crop. A higher temperature should be maintained during the photoperiod, while a fall in temperature during the dark period can aid in stress recovery. At UQ, a 12 hour 22 °C / 17 °C temperature cycling regime with the 2 hours of darkness occurring within the 12 hours of 17 °C has proven successful (Speed breeding II)^1^. In contrast, a temperature cycling regime of 22 °C / 17 °C for 22 hours light and 2 hours dark, respectively, is used at JIC (Speed breeding I)^1^. In both scenarios, the generation times of all crops were successfully accelerated and comparable. In the controlled environment chamber in which this was demonstrated, the temperature was ramped up and down similarly to the lights, but this was subsequently found to not be of particular importance.

##### iv) Humidity

Most controlled environment chambers have limited control over humidity but a reasonable range of 60-70% is ideal. For crops that are more adapted to drier conditions, a lower humidity level may be advisable.

#### Section b) Benchtop growth cabinet

To construct your low cost growth cabinet the following components are required.

##### Hardware

- 12 V, 50 A DC power supply 600 W (Amazon, cat. no. B072M7P7QJ)
- 12 V to 5 V, 3 A DC/DC converter module (Amazon, cat. no. B00G890MIC)
- USB extension cable – 30 cm (Amazon, cat. no. B002M8RVKA)
- Ethernet extension cable – 30 cm (Amazon, cat. no. B077V421QH)
- Arduino UNO (Amazon, cat. no. B00CGU1VOG)
- Raspberry Pi 3 model B (CPC, cat. no. 2525225)
- Raspberry Pi display 7 inch touchscreen (CPC, cat. no. 2473872)
- Arduino base shield v2 – SeeedStudio (CPC, cat. no. SC13822)

##### Cabinet structure

- Aluminium composite panel, 757 × 307 × 3 mm, quantity = 6 (Cut Plastics, cat. no. CP027-03)
- Aluminium composite panel, 757 × 357 × 3 mm (Cut Plastics, cat. no. CP027-03)
- Aluminium composite panel, 757 × 107 × 3 mm (Cut Plastics, cat. no. CP027-03)
- Aluminium composite panel, 757 × 757 × 3 mm (Cut Plastics, cat. no. CP027-03)
- PVC foam board, 757 × 157 × 3 mm, quantity = 2 (Cut Plastics, cat. no. CP015-03)
- PVC foam board, 757 × 141 × 3 mm (Cut Plastics, cat. no. CP015-03)
- PVC foam board, 757 × 307 × 3 mm, quantity = 2 (Cut Plastics, cat. no. CP015-03)
- Perspex clear acrylic sheet, 757 × 307 × 3 mm (Cut Plastics, cat. no. CP001-03)
- OpenBeam, 1000 mm, quantity = 4 (Technobots Online, cat. no. 4451-900)
- OpenBeam, 750 mm, quantity = 13 (Technobots Online, cat. no. 4451-750)
- OpenBeam, 300 mm, quantity = 10 (Technobots Online, cat. no. 4451-300)
- Corner bracket – MakerBeam, quantity = 4 (Technobots Online, cat. no. 4446-013)
- L-joining plate – OpenBeam, quantity = 36 (Technobots Online, cat. no. 4450-003)
- T-joining plate – OpenBeam, quantity = 2 (Technobots Online, cat. no. 4450-004)

##### Lighting system

- Full spectrum grow light LED bulb, quantity = 16 (Amazon, cat. no. 071J3BC1W)
- E27 lamp holder, quantity = 16 (Sinolec Components, cat. no. E27-SD04-2)
- Solid state relay – grove SeedStudio (Mouser, cat. no. 713-103020004)

##### Temperature and humidity control system

- 12 V, 10 A thermoelectric cooler, quantity = 3 (Amazon, cat. no. B01M2ZBBVM)
- Temperature and humidity sensor pro–grove SeeedStudio (CPC, cat. no. MK00343)
- Relay – grove SeedStudio, quantity = 4 (CPC, cat. no. MK00330)
- 12 V cooling fan, 50 mm (Amazon, cat. no. B00HPKC5MO) Software
- Arduino IDE (v1.8.5, https://www.arduino.cc/en/Main/Software)

#### Section c) LED-supplemented glasshouse setup

##### i. Glasshouse

A well-located glasshouse with the required space and sufficient ambient lighting. We recommend fitting a temperature control system and programmable lights. Controllable blinds are also optional if blocking out high irradiance on very sunny days is required.

##### ii. LED lamps

While any kind of lighting system can be used to supplement the ambient lighting in the glasshouse, we recommend LED lamps above all because of the significant savings these provide in terms of maintenance and energy consumption. The glasshouse-based SB experiments detailed in our previous paper^1^ were based on SVLs, but we have obtained similar results with LED-lighting at both UQ and JIC. The lighting system configuration, make and model of the lights for both locations are provided in Equipment setup.

##### iii. SSD trays

For demonstration, at UQ, three seedling tray types with increasing sowing densities were used. The dimensions and volumes are given in Supplementary Table 7. The soil media composition is given in Supplementary Table 4.

###### Caution

Energy tariffs can vary according to the time of day, depending on peak energy usage patterns in the location. Substantial savings can be achieved by programming the dark period to coincide with the energy tariff imposed during peak electricity consumption.

#### Additional equipment needed

##### i. PAR meter

The PAR is measured in either PPFD or Lux. Any off-the-shelf PAR meter can be used, as long as it provides PPFD levels and relative wavelength composition. We used the MK350S Spectrometer from UPRtek and the Spectrum Genius Essence Lighting Passport light sensor from AsenseTek Inc. (Taiwan) at JIC and UQ, respectively.

##### ii. Energy meter

This allows measuring the energy consumption for lighting and temperature maintenance thereby providing insight into SB operational costs. Any off-the-shelf energy meter can be used for this purpose. To obtain energy consumption data for both the lights employed and the Controlled Environment Rooms (CERs) at JIC, we utilised a clamp-on Current Transformer meter with the capacity to store and download data. The instrument provided half hourly readings and as such was highly accurate in determining energy costs

### Equipment setup

In this section, we provide detailed protocols for the SB setups discussed in the previous section.

#### a) Benchtop growth cabinet

Hardware: Connect the display to the Raspberry Pi using the provided cables as instructed by the manufacturer. The Arduino connects to the Raspberry Pi via USB ports. Sensors and relay modules are connected using the Grove system (SeedStudio).
Cabinet structure: Assemble the beam profile using the joining plates. Slide the panels, boards and sheets before fully assembling each side.
Lighting system: The photoperiod with the full-spectrum LED light bulbs is controlled by a solid-state relay connected to the Arduino microcontroller. Sixteen 57 mm diameter holes need to be drilled in one of the 757 × 307 × 3 mm aluminium composite panels, to fit the E27 lamp holders. The lamp holders are then inserted and wired in parallel.
Temperature and humidity system: Pre-assembled thermoelectric cooling modules are used to simplify the construction of the benchtop growth cabinet. These are composed of fans, aluminium heat sinks, and Peltier elements. The cooling modules are controlled by relays connected to the Arduino. Airflow is used to control the humidity, *i.e.* the humidity sensor will trigger the 12 V fan to circulate air from outside the cabinet in order to reduce the humidity inside.
Software installation and setup: The speed breeding cabinet is controlled by three main subsystems: The arduino micro controller that monitors and controls the environment according to a desired optimal; a python daemon that stores the current conditions and reads the expected conditions from a MongoDB database and; a graphical interface written in ReactJS that allows the users to set up the expected conditions in a 24-hour range.

The circuit diagram for making the connections are provided in Supplementary Figure 3 and a photograph of the assembled cabinet is provided in Supplementary Figure 4. The cabinet has an available area of 0.225 m^2^. For the lamps we have used, the spectrum is provided in Supplementary Figure 5, with the light levels in PPFD (Photosynthetic Photon Flux Density) being on an average about 120 μmol·m^-2.^s^-1^ at 16 cm above the base where the pots are kept, and about 320 μmol·m^-2.^s^-1^ and 220 μmol·m^-2.^s^-1^ from a 10 cm and 20 cm distance respectively from the top of the cabinet where the lights are situated. The energy consumption of the mini cabinet is 6.24 kWh per day.

##### NOTE

A step-by-step guide for constructing the cabinet and installing the software is available at https://github.com/PhenoTIPI/SpeedSeed3/wiki, along with troubleshooting tips.

##### Caution

The construction of the cabinet requires the use of sharp cutting and drilling tools that may cause physical injury if handled improperly. Many steps involve electrical components, which can cause fire if operated without being earthed. Ensure all necessary safety steps are followed and use personal protective equipment when constructing the cabinet.

#### b) LED-supplemented glasshouse

<u>Table 2</u> provides the lighting arrangement in two glasshouse configurations. Both setups have been demonstrated to successfully support SB for the species listed.

**Table 2.**
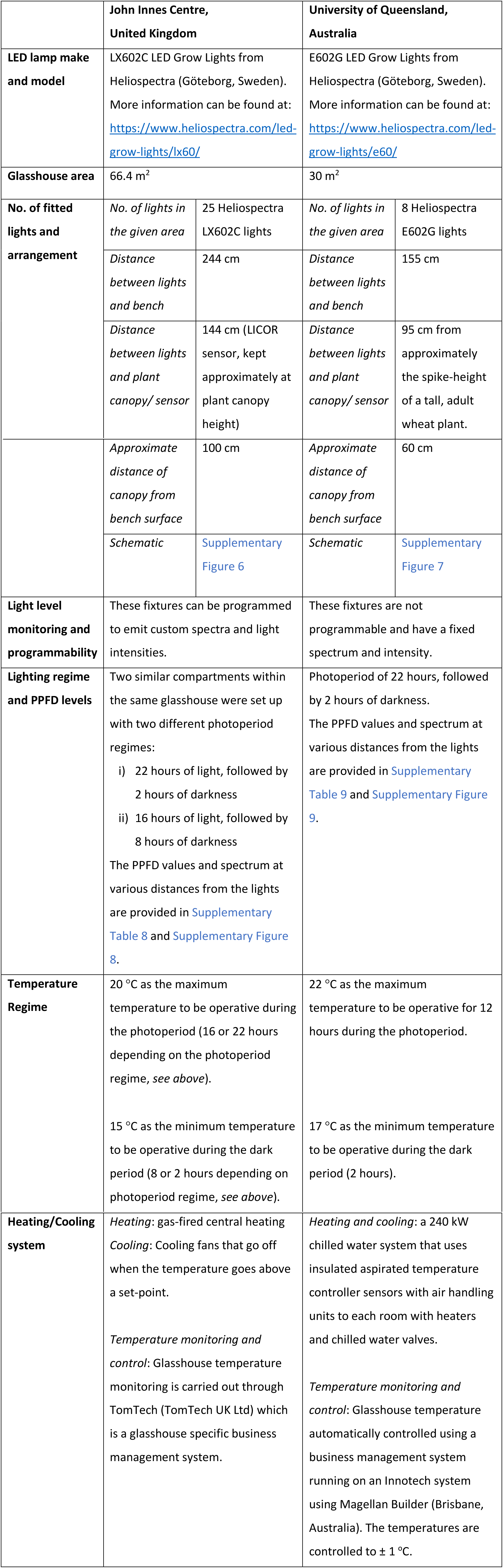
LED-Supplemented Glasshouse setups for speed breeding at JIC and UQ.

|  | John Innes Centre,<br>United Kingdom |  | University of Queensland,<br>Australia |  |
| --- | --- | --- | --- | --- |
| **LED lamp make and model** | LX602C LED Grow Lights from Heliospectra (Göteborg, Sweden).<br>More information can be found at: https://www.heliospectra.com/led-grow-lights/lx60/ |  | E602G LED Grow Lights from Heliospectra (Göteborg, Sweden).<br>More information can be found at: https://www.heliospectra.com/led-grow-lights/e60/ |  |
| **Glasshouse area** | 66.4 m<sup>2</sup> |  | 30 m<sup>2</sup> |  |
| **No. of fitted lights and arrangement** | *No. of lights in the given area* | 25 Heliospectra LX602C lights | *No. of lights in the given area* | 8 Heliospectra E602G lights |
|  | *Distance between lights and bench* | 244 cm | *Distance between lights and bench* | 155 cm |
|  | *Distance between lights and plant canopy/ sensor* | 144 cm (LICOR sensor, kept approximately at plant canopy height) | *Distance between lights and plant canopy/ sensor* | 95 cm from approximately the spike-height of a tall, adult wheat plant. |
|  | *Approximate distance of canopy from bench surface* | 100 cm | *Approximate distance of canopy from bench surface* | 60 cm |
|  | *Schematic* | Supplementary Figure 6 | *Schematic* | Supplementary Figure 7 |
| **Light level monitoring and programmability** | These fixtures can be programmed to emit custom spectra and light intensities. |  | These fixtures are not programmable and have a fixed spectrum and intensity. |  |
| **Lighting regime and PPFD levels** | Two similar compartments within the same glasshouse were set up with two different photoperiod regimes:<br>i) 22 hours of light, followed by 2 hours of darkness<br>ii) 16 hours of light, followed by 8 hours of darkness<br>The PPFD values and spectrum at various distances from the lights are provided in Supplementary Table 8 and Supplementary Figure 8. |  | Photoperiod of 22 hours, followed by 2 hours of darkness.<br>The PPFD values and spectrum at various distances from the lights are provided in Supplementary Table 9 and Supplementary Figure 9. |  |
| **Temperature Regime** | 20 °C as the maximum temperature to be operative during the photoperiod (16 or 22 hours depending on the photoperiod regime, *see above*).<br>15 °C as the minimum temperature to be operative during the dark period (8 or 2 hours depending on photoperiod regime, *see above*). |  | 22 °C as the maximum temperature to be operative for 12 hours during the photoperiod.<br>17 °C as the minimum temperature to be operative during the dark period (2 hours). |  |

|  |  |  |
| --- | --- | --- |
| **Heating/Cooling system** | *Heating:* gas-fired central heating<br>*Cooling:* Cooling fans that go off when the temperature goes above a set-point.<br>*Temperature monitoring and control:* Glasshouse temperature monitoring is carried out through TomTech (TomTech UK Ltd) which is a glasshouse specific business management system. | *Heating and cooling:* a 240 kW chilled water system that uses insulated aspirated temperature controller sensors with air handling units to each room with heaters and chilled water valves.<br>*Temperature monitoring and control:* Glasshouse temperature automatically controlled using a business management system running on an Innotech system using Magellan Builder (Brisbane, Australia). The temperatures are controlled to $\pm 1$ °C. |

##### Critical

Weather and ambient light varies by location and season, especially at higher latitudes. Thus, for the glasshouse setups listed here, the light spectrum is determined not just by the presence of the LED lights but also by the ambient light. To ensure reproducibility, consider setting up your experiment in a way that mitigates these environmental variables. For example, use programmable lights that allow intensity modification based on sensor feedback, or controllable blinds to regulate photoperiod. Provision of a short dark-period is recommended for optimum plant health. We highly recommend setting up a temperature monitoring and control system.

A summary of the crops for which we have successfully demonstrated a shortening of generation time using SB, including information on which specific SB setups were used, and where the reader can find more information on the key growth stages and other growth parameters of the crop grown under those conditions is provided in <u>Table 3</u>.

**Table 3.** A list of speed breeding protocols that have been demonstrated for different species along with pointers for locating the associated data.

| Species | Demonstrated SB conditions and associated data |  |  |
| --- | --- | --- | --- |
|  | This protocol | Watson and Ghosh et al., 2018 | Other |
| Spring wheat<br>*T. aestivum* | JIC-GH-LED<sup>1</sup><br>(Supplementary Tables 10 - 18)<br>UQ-GH-LED<sup>2</sup><br>(Supplementary Tables 19 and 20) | UQ-GH-SVL<sup>3</sup><br>(Supplementary Tables 11, 15, 21, 28, 30, 31)<br>CER-JIC<sup>4</sup><br>(Supplementary Tables 2, 5-8, 19, 27, 34-36) |  |
| Winter wheat<br>*T. aestivum* | JIC-GH-LED<br>(Supplementary Tables 21 - 23) |  |  |
| Durum wheat<br>*T. durum* | JIC-GH-LED<br>(Supplementary Tables 14 - 18) |  | Alahmad et al., 2018 |
| Spring barley<br>*H. vulgare* | JIC-GH-LED<br>(Supplementary Tables 24 - 26)<br>UQ-GH-LED<br>(Supplementary Tables 27 and 28) | UQ-GH-SVL<br>(Supplementary Tables 12, 16, 20, 22, 29, 30, 32)<br>CER-JIC<br>(Supplementary Tables 3, 6, 37, 38) |  |
| Canola<br>*Brassica napus* | JIC-GH-LED<br>(Supplementary Tables 29 - 33) | UQ-GH-SVL<br>(Supplementary Tables 13, 17, 23, 25, 30, 39) |  |
| *Brassica rapa* | JIC-GH-LED<br>(Supplementary Tables 29 - 33) |  |  |
| *Brassica oleracea* | JIC-GH-LED<br>(Supplementary Tables 29 - 33) |  |  |
| Pea | JIC-GH-LED | CER-JIC |  |
| *P. sativum* | (Supplementary Tables 34 and 35) | (Supplementary Table 10) |  |
| Grasspea<br>*L. sativus* | JIC-GH-LED<br>(Supplementary Tables 36 - 38) |  |  |
| Medicago |  | CER-JIC<br>(Supplementary Table 9) |  |
| *Brachypodium distachyon* | JIC-GH-LED<br>(Supplementary Tables 39, 40) | CER-JIC<br>(Supplementary Table 4) |  |
| Quinoa<br>*C. quinoa* | JIC-GH-LED<br>(Supplementary Tables 41 - 43) |  |  |
| Oat<br>*A. strigosa* | JIC-GH-LED<br>(Supplementary Tables 44 - 46) |  |  |
| Chickpea<br>*C. arietinum* |  | UQ-GH-SVL<br>(Supplementary Tables 14, 18, 24, 26, 30) |  |
| Peanut<br>*A. hypogaea* |  |  | O'Connor et al., 2013<sup>10</sup> |
| Amaranth<br>*Amaranthus* spp. |  |  | Stetter et al., 2016<sup>36</sup> |

<sup>1</sup> **JIC-GH-LED**: LED-Supplemented Glasshouse setup, JIC, UK (described in this paper, Equipment Setup Section c).
<sup>2</sup> **UQ-GH-LED**: LED-Supplemented Glasshouse setup, UQ, Australia (described in this paper, Equipment Setup Section c).
<sup>3</sup> **UQ-GH-SVL**: SVL-Supplemented Glasshouse setup, UQ, Australia (described in Methods section: Speed Breeding II<sup>1</sup>).
<sup>4</sup> **CER-JIC**: Controlled Environment Room, JIC, UK (described in Methods section: Speed Breeding I<sup>1</sup>).

## 4. Procedure

### a) Preparing seed for sowing

To increase germination efficiency some seeds may need a pre-treatment either by cold stratification (prolonged imbibition in the cold) or scarification (physical or chemical weakening of the seed coat). The requirements for germination pre-treatments are specific for each species, and accessions of that species, and should be determined on an individual basis. Dormant spring wheat and barley seed can be imbibed on moistened filter paper in a Petri dish for 24 hours and then chilled at 4 °C for approximately three days (longer times may be required depending on the level of dormancy). The seeds can then be left at room temperature for one to three days to germinate prior to transferring to soil. If pre-treatment is not required, the seed can be germinated in a Petri dish on moistened filter paper before transferring to soil. In a large-scale scenario, seeds can be directly sown into high density trays and placed in a cold-room, then trays can be moved to the growing environment in the glasshouse. If a pre-treatment is not required, seed may be sown directly into soil in the glasshouse/growth chamber.

#### Caution

If seeds germinate in a Petri dish and become too well established (i.e. develop green leaves) before transplanting to soil, the shift to SB conditions, especially the presence of intense light, can shock the plants, resulting in a strong hypersensitive response and possibly death. Take care to prick them out early, or if they are already established, transfer them to soil and place a mesh over the plants to reduce light intensity while they adapt to the new environmental conditions.

### b) Monitoring key growth stages, growth parameters, and phenotyping

To enable comparison to normal development, monitor the key growth stages of the plants. For many crops, defined growth stages have been published; for example, cereal crops^40^, canola^41^, quinoa^42^ and legumes^43^. Take note of the heading times and earliest time point to harvest viable seeds. We also advise monitoring the height and general physiology of the plants.

#### NOTE

Experiments performed in Section c, LED-supplemented glasshouse setup at the JIC, UK, involved a SB glasshouse compartment as detailed above (i.e. 22 h day length), and a twin compartment with a 16 h day length to measure the effect and value of increased day length. Growth parameters and harvest times are provided for both lighting regimes where available.

For wheat and barley, we have also previously demonstrated how SB conditions do not interfere with the phenotyping of a number of key traits^1^, and how variations of the SB protocol can be used to rapidly screen wheat and barley for resistance to a number of major diseases or disorders (<u>Table 4</u>).

**Table 4.** Protocols for phenotyping diseases and disorders under speed breeding conditions.

| Disease / disorder | Species | Reference |
| --- | --- | --- |
| Stripe rust<br>(*Puccinia striiformis* f. sp. *tritici*) | Spring wheat<br>(*T. aestivum*) | Pretorius *et al.* (2000). *Acta Phytopathologica et Entomologica Hungarica*, 35(1-4), 359-364<sup>44</sup><br>Hickey *et al.* (2012). *Plant Breeding*, 131(1), 54-61<sup>14</sup> |
| Leaf rust<br>(*Puccinia recondita* f. sp. *tritici*, “brown rust”)<br>(*Puccinia triticina*, “black rust”) | Spring wheat<br>(*T. aestivum*) | Pretorius *et al.* (2000). *Acta Phytopathologica et Entomologica Hungarica*, 35(1-4), 359-364<sup>44</sup><br>Riaz *et al.* (2016). *Plant Methods*, 12, 17<sup>14</sup> |
| Yellow spot / Tan spot<br>(*Pyrenophora tritici-repentis*) | Spring wheat<br>(*T. aestivum*) | Dinglasan *et al.* (2016). *Euphytica*, 209(3), 693-707<sup>12</sup> |
| Leaf rust<br>(*Puccinia hordei*)<br>Net form net blotch<br>(*Pyrenophora teres* f. sp. *teres*)<br>Spot form net blotch<br>(*Puccinia teres* f. sp. *maculate*)<br>Spot blotch<br>(*Cochliobolus sativus*) | Barley<br>(*H. vulgare*) | Hickey *et al.* (2017). *Euphytica*, 213(3), 64<sup>15</sup> |
| Stem rust<br>(*Puccinia graminis* f. sp. *tritici*) | Spring wheat<br>(*T. aestivum*) | Riaz and Hickey (2017). *Wheat Rust Diseases: Methods and Protocols* (Vol. 1659, pp. 183-196)<sup>45</sup> |
| Crown rot<br>(*Fusarium pseudograminearum*) | Durum wheat<br>(*T. durum*) | Alahmad *et al.* (2018). *Plant Methods*, 14(1), 36<sup>11</sup> |
| Pre-harvest sprouting | Spring wheat<br>(*T. aestivum*) | Hickey *et al.* (2009). *Euphytica* 168, 303-310<sup>9</sup> |
| Pod shattering | Canola<br>(*B. napus*) | Watson and Ghosh *et al.* (2018). *Nature Plants*, 4(1), 23-29<sup>1</sup> |

### c) Seed harvesting

Shortened generation times can also be achieved in some species by harvesting premature seed. This usually involves waiting until the seeds have set in the plant (indicated by filled seed in spikes for wheat, or filled pods for legumes), then either increasing the temperature or withholding water from the plant to hasten seed ripening and drying. After a week of this stress application, seeds may be harvested.

#### NOTE

For experiments performed in the third protocol setup (Section c, LED-supplemented glasshouse) at the JIC, UK, early harvest times are provided for both lighting regimes where available. If not indicated, the harvest time outlined is for harvest at physiological maturity.

#### Caution

Freshly harvested seed may display dormancy. See troubleshooting (Section 5) for more details on how to overcome this issue.

### d) Monitoring energy use

At the end of one cycle, review the energy costs for your SB system. This is particularly useful to evaluate the generation time vs cost trade-off where multiple conditions have been tested concurrently (e.g. different day lengths). For the LED-Supplemented glasshouse setup in JIC, there were two rooms set up concurrently with 16-hour and 22-hour photoperiods. The energy calculations for running each of these setups per month is given in Supplementary Table 47, along with a comparison of how much it would cost to run a similar setup with Sodium Vapour Lamps.

## 5. Troubleshooting

**Table 5.** Suggested solutions to common issues under speed breeding.

| Problem | Possible Reason | Solution |
| --- | --- | --- |
| Plants exhibit tip-burn necrosis. The leaves curl inward or outward, and may have small, circular depressions or “bubbles” (Supplementary Figure 2). | Calcium deficiency – common in accelerated growth. | Apply a liquid fertiliser containing calcium as a foliar spray early in growth to control any developing deficiency. This may be a 1% calcium nitrate solution applied 2-3 times per week or as part of another broad-spectrum fertiliser.<br>Acidic soil can interfere with calcium uptake – adding dolomite to the soil can reduce acidity if the base soil mix tends to a lower pH. |
| Initially curling and death of young leaf-tips and down the leaf blade. Young leaves may also not emerge properly and form loops or twists. Later, spike top can wither, turn white and fail to produce grain. Spikes may also become twisted into curls (Supplementary Figure 10). | Copper deficiency – common in accelerated growth. | Apply a liquid fertiliser containing copper as a foliar spray early in growth to control any developing deficiency.<br>Alkaline or waterlogged soil can affect copper uptake – do not over-water or add excessive dolomite when ameliorating calcium deficiency as described above. |
| Young leaves appear striped with interveinal yellowing (Supplementary Figure 11). | Iron deficiency. | Apply a liquid fertiliser containing iron as a foliar spray early in growth to |
|  |  | control any developing deficiency. |
| Plants are weak and spindly or suffering chlorosis. | These are possible symptoms of a range of nutrient deficiencies. | Apply a liquid fertiliser with a broad range of nutrients to the soil and as a foliar spray. |
| Seeds do not germinate. | Seed harvested too early and are not viable.<br>Seeds are dormant. | Harvest seed slightly later.<br>Store the seeds for a few additional days or weeks before trying again.<br>Alternatively, cold stratify the seed at 4-5 °C for several days and/or treat with a low concentration (~0.5 ppm) of gibberellic acid (GA3) by dipping the seeds into the solution or spraying. |
| Plants did not cycle much faster than in the glasshouse with no supplemental lights and/or in field conditions, even though they are LD or day neutral plants. | The optimum conditions for rapid generation advancement have not been reached for the crop.<br>The particular genotype may be recalcitrant to SB. | Make adjustments for temperature, light intensity, light quality and/or day length.<br>Try other genotypes to explore if it is a genotype- or species-specific issue. |
| LD or day neutral plants do not flower. | Vernalisation needed. | Depending on the species, vernalise the plants for up to 8 weeks at 4 to 10 °C. |

## 6. Anticipated Results

As demonstrated in our previous study, under SB conditions with a 22-hour photoperiod, it should be possible to produce up to 6 generations per year in spring wheat and barley and up to 4 and 4.5 generations per year in canola and chickpea, respectively^1^. However, it is important to remember that results are highly dependent on the crop species and can vary greatly between cultivars. The light quality, duration of the photoperiod and temperature regime also impact the extent to which the generation time is reduced. It should also be noted that ambient sunlight strength and duration will vary with location and season, thus resulting in differences in rate of development. These factors, in addition to basic growing conditions, such as soil type, can be manipulated to obtain the optimal parameters for the crop of interest. The various protocols outlined above are designed to facilitate this process.

The self-made, bench-top speed breeding cabinet will facilitate identification of conditions that enable rapid-cycling of wheat and pea, and by extension, the other crops listed (Supplementary Figure 4). We demonstrated the efficacy of this cabinet design by growing rapid-cycling varieties of pea (*P. sativum* cv. JI 2822) and wheat (*T. aestivum* cv. USU Apogee) and showing the shortened time from seed to seed, without compromising the viability of early harvested seed (Supplementary Tables 1, 2). This is comparable with data from our previous study^1^ where we evaluated the same pea variety (JI 2822) under SB conditions using a commercial CER.

The time taken for reproductive development to occur for a range of crop species under the LED-fitted, SB glasshouse (JIC, UK) is provided in <u>Table 6</u>. Two extended photoperiods are represented to give an approximate expectation of the rapid development of these species under SB, and to give the reader an idea of what a 6-hour difference in photoperiod can produce in a range of crops and cultivars. The much slower rate of development under control or regular glasshouse conditions without supplemental lighting was reported for some of these species in our previous study^1^. Plants grown under SB can be expected to look healthy (<u>Figure 1</u>) with minor reductions in seed set (refer to <u>Table 3</u> in order to view the related data for the crop of interest) and spike size (Supplementary Figure 12) or pod size (Supplementary Figures 13 and 14). In some crop species, the SB conditions can produce a slight reduction in height and/or internode length. In our experience, while working on *M. truncatula* and *P. sativum*, we found the plants grown under SB produced leaves with much smaller surface areas. Occasionally, micronutrient deficiencies manifest themselves because of the rapid growth and change in soil pH – some of these issues (particularly for wheat and barley) are highlighted in the Troubleshooting section. Despite efforts to optimise soil composition, there may be a cultivar that responds very poorly to the long-photoperiod and high irradiance.

**Figure 1.**
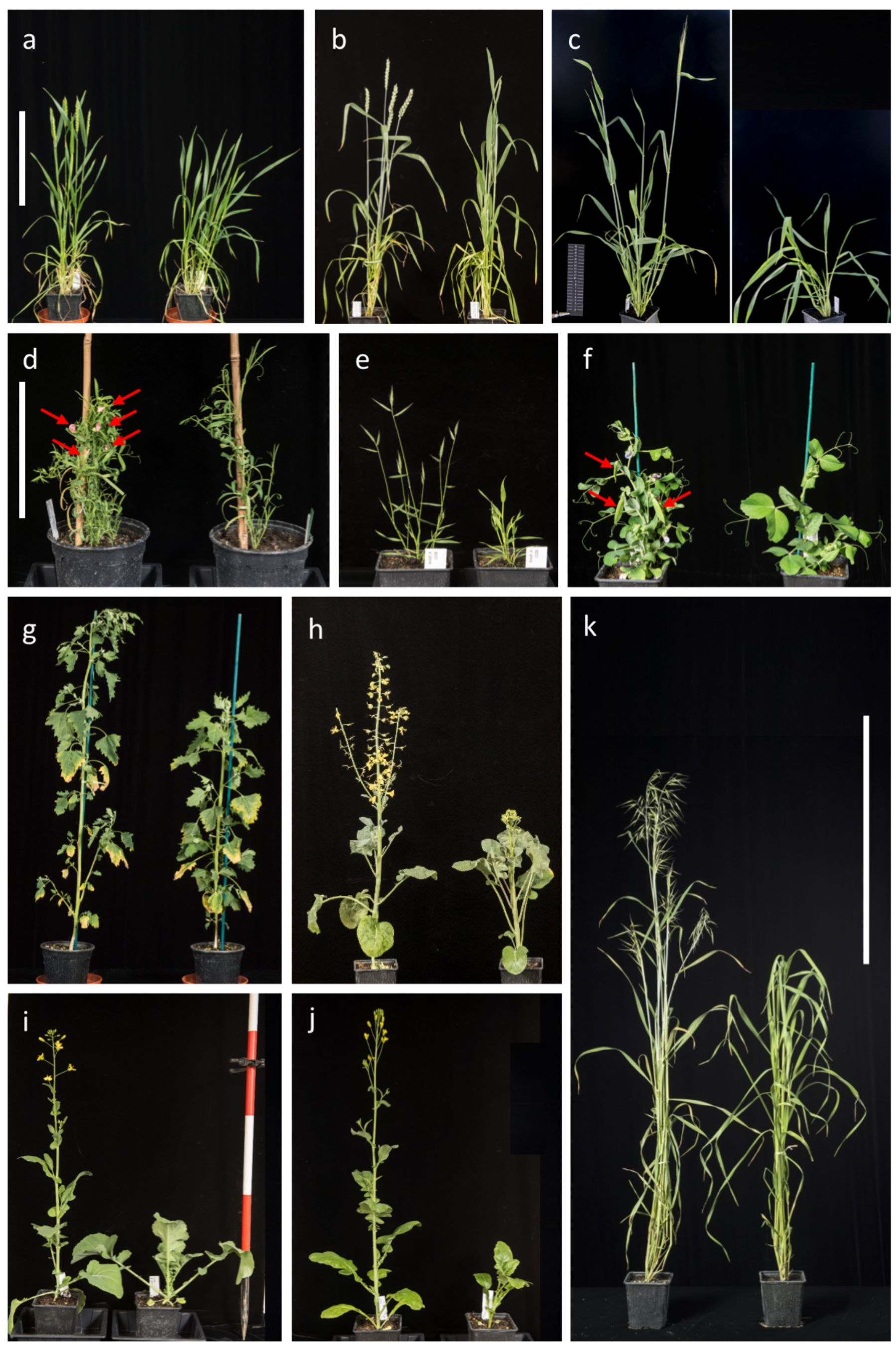
Accelerated plant growth and development under speed breeding (22 hour photoperiod conditions) (left) compared to standard long-day conditions (16 hour photoperiod) (right) in LED-supplemented glasshouses at John Innes Centre, UK. a, Winter growth-habit wheat (*T. aestivum* cv. Crusoe) at 112 days after sowing (DAS), including 12 days of growth under 16-hour photoperiod conditions followed by 56 days of vernalisation at 6 °C with 8 hour photoperiod; **b,** Spring wheat (*T. aestivum* cv. Cadenza) at 57 DAS; **c,** Spring barley (*H. vulgare* cv. Manchuria) at 35 DAS; (scalebar is 20 cm for a, b, c) **d,** Grasspea (*L. sativus* cv. Mahateora) at 35 DAS (red arrows indicate position of flowers); **e,** *B. distachyon* (accession Bd21) at 34 DAS; **f,** Pea (*P. sativum* accession JI 2822) at 34 DAS; (scalebar is 20 cm for d, e, f) **g,** Quinoa (*C. quinoa* accession QQ74) at 58 DAS; **h,** *Brassica oleracea* (line DH1012) at 108 DAS; **i,** *Brassica napus* (line RV31) at 87 DAS; **j,** *Brassica rapa* (line R-0-18 87) at 87 DAS; **k,** Diploid Oat (*A. strigosa* accession S75) at 52 DAS (scalebar is 60 cm for g, h, i, j). All plants were sown in October or November 2017, except for the quinoa, which was sown in February 2018.

We have previously demonstrated that wheat, barley and canola plants grown under SB are suitable for crossing and phenotyping a range of adult plant traits^1^. That said, complex phenotypes such as yield and abiotic stress resilience (heat or drought stress) are best evaluated in the field, particularly for breeding objectives. We have also demonstrated how SB can be combined with transformation of barley to speed up the process of obtaining transformed seeds^1^.

**Table 6.** Mean days to anthesis^1^ under speed breeding using LED-supplemented glasshouses at JIC, UK. All plants had a temperature cycle regime of 22 hours at 22 °C and 2 hours at 17 °C to coincide with the light and dark period, respectively.

| Species | Associated data | Photoperiod | Mean days to flowering<sup>1</sup> |
| --- | --- | --- | --- |
| Spring wheat<br>*T. aestivum* | Supplementary Tables 10-18, 19, 20 | 22 h | 49.6 ± 5.0 |
|  |  | 16 h | 62.5 ± 4.3 |
| Winter wheat<br>*T. aestivum* | Supplementary Tables 21 - 23 | 22 h | 105.4 ± 1.7 |
|  |  | 16 h | 115.4 ± 1.9 |
| Durum wheat<br>*T. durum* | Supplementary Tables 14 - 18 | 22 h | 46 ± 1.9<sup>2</sup> |
|  |  | 16 h | 53.7 ± 1.0<sup>2</sup> |
| Spring barley<br>*H. vulgare* | Supplementary Tables 24 - 28 | 22 h | 38.4 ± 13.9 |
|  |  | 16 h | 46.6 ± 12.1 |
| Canola<br>*Brassica napus* | Supplementary Tables 29 - 33 | 22 h | 34.5 ± 0.7<sup>3</sup> |
|  |  | 16 h | 45.0 ± 0.0 |
| *Brassica rapa* | Supplementary Tables 29 - 33 | 22 h | 36.5 ± 2.5<sup>3</sup> |
|  |  | 16 h | 41.0 ± 3.7 |
| *Brassica oleracea* | Supplementary Tables 29 - 33 | 22 h | 49.2 ± 1.8<sup>3</sup> |
|  |  | 16 h | 61.2 ± 2.3 |
| Pea<br>*P. sativum* | Supplementary Tables 34, 35 | 22 h | 32.2 ± 5.3<sup>4</sup> |
|  |  | 16 h | 42.9 ± 5.3 |
| Grasspea<br>*L. sativus* | Supplementary Tables 36 - 38 | 22 h | 31<sup>3</sup> ± |
|  |  | 16 h | ND |
| *Brachypodium distachyon* | Supplementary Tables 39, 40 | 22 h | 31.5 ± 5.2 |
|  |  | 16 h | 44.0 ± 5.2 |
| Quinoa<br>*C. quinoa* | Supplementary Tables 41 - 43 | 22 h | 54.6<sup>5</sup> ± 0.6 |
|  |  | 16 h | 61.1 ± 4.6 |
| Oat | Supplementary Tables 44 - 46 | 22 h | 52 ± 0.0 |
| *A. sativa* |  | 16 h | 66 ± 0.0 |

<sup>1</sup>Days to flowering/anthesis (GS65) from sowing<sup>40</sup>.
<sup>2</sup>Days to 50% ear emergence from sowing (GS55).
<sup>3</sup>Days to first flower opening from sowing.
<sup>4</sup>Days to the first flower bud from sowing.
<sup>5</sup>Days to anthesis (growth stage 6 according to BBCH scale<sup>42</sup>).

In breeding programs, SSD is often an important step in cultivar development that requires high-density plantings. The SB protocols provided for glasshouses are ideal for SSD programs, particularly cereal crops. Increasing sowing density under SB can enable rapid cycling of many lines with healthy plants and viable seed. <u>Figure 2</u> shows an example of the plant condition, spike lengths and seed sizes that could be expected at various sowing densities in SB. Under the UQ-GH-LED protocol, at a density of 1000 plants/m^2^, up to 6 generations of wheat and barley can be expected per year (Supplementary Table D and E). At higher densities, plant height and seed numbers can be reduced due to the greater competition and low soil volume. Despite this, even at the highest sowing density shown here, all plants produced a spike with at least enough seed to perform SSD, and in most cases many more. Large differences in the speed of development can be achieved by extending the photoperiod from 16 to 22 hours. Under the JIC-GH-LED protocol, spring and durum wheat were over ten days faster in development with an additional 6 hours of photoperiod. <u>Table 7</u> provides the approximate development times for several cereal crops at a range of sowing densities, appropriate for intensive SSD. The SSD SB protocol was performed under two extended photoperiod and temperature regimes at either JIC, UK, or UQ, Australia. These results demonstrate that plants can be grown at high densities under SB conditions to produce plants suitable for effective and resource-efficient generation turnover in SSD programs.

**Figure 2.**
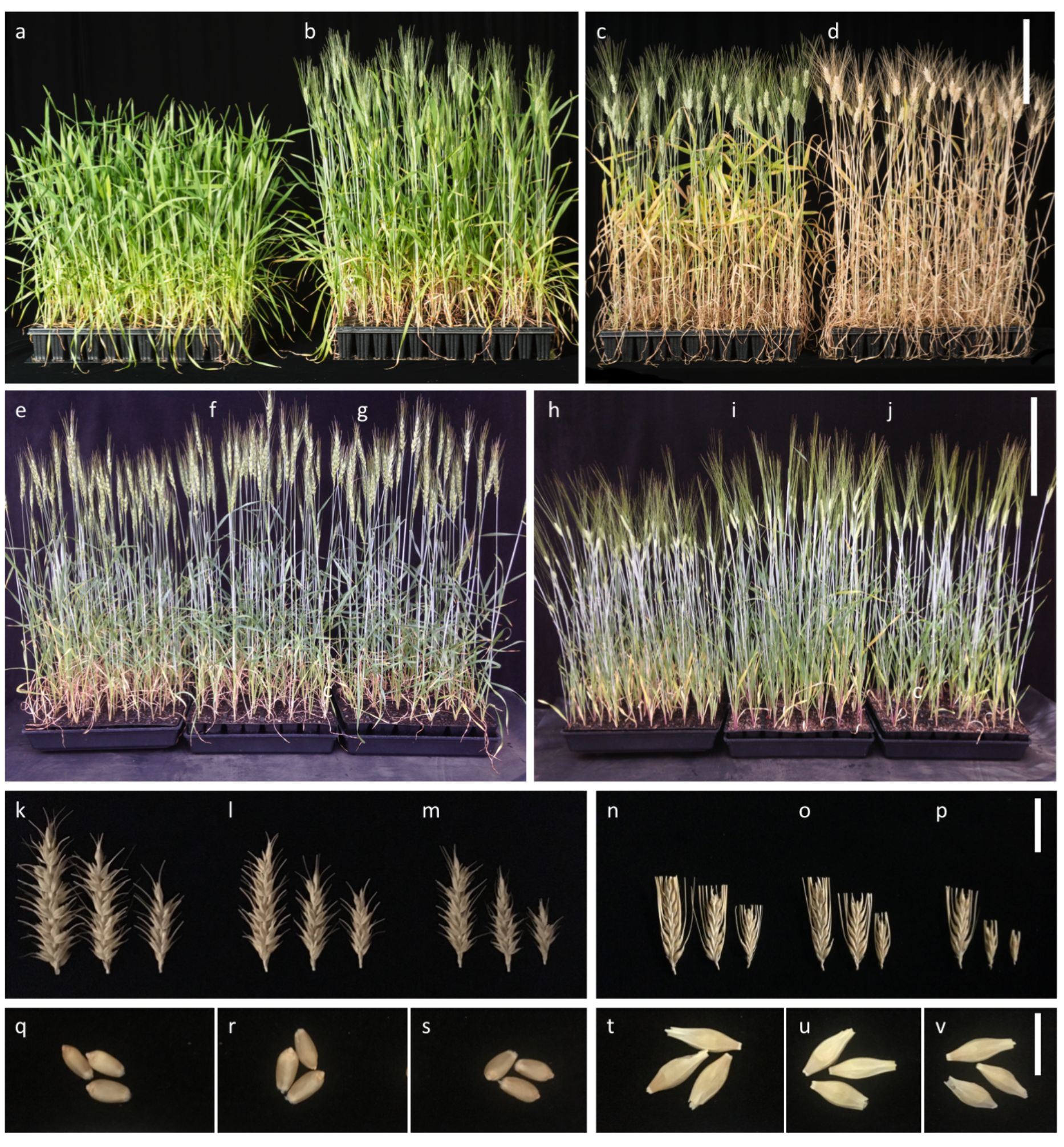
Single seed descent sowing densities of wheat (spring and durum) and barley under LED-Supplemented Glasshouse setup at JIC, UK and UQ, Australia. Durum wheat (*T. durum* cv. Kronos) grown under the LED-Supplemented Glasshouse setup, JIC, UK, in 96-cell trays: **a**, Forty-three days after sowing under 16-hour photoperiod; **b**, Forty-three days after sowing under 22-hour photoperiod; **c**, Seventy-nine days under 16-hour photoperiod; **d**, Seventy-nine days under 22-hour photoperiod. Scale bar is 20 cm. Spring wheat (*T. aestivum* cv. Suntop) grown under LED-Supplemented Glasshouse setup, UQ, Australia, at 37 days after sowing: **e**, plants in a 30-cell tray; **f**, plants in a 64-cell tray; **g**, plants in a 100-cell. Barley (*H. vulgare* cv. Commander) grown under LED-Supplemented glasshouse setup, UQ, Australia, at 34 days after sowing: **h**, plants in a 30-cell tray; **i**, plants in a 64-cell tray; **j**, plants in a 100-cell. Scale bar is 20 cm. Mature spikes of spring wheat (*T. aestivum* cv. Suntop) grown under LED-Supplemented glasshouse setup, UQ, Australia: **k**, plants in a 30-cell tray; **l**, plants in a 64-cell tray; **m**, plants in a 100-cell. Mature spikes of barley (*H. vulgare* cv. Commander) grown under LED-Supplemented glasshouse setup, UQ, Australia: **n**, plants in a 30-cell tray; **o**, plants in a 64-cell tray; **p**, plants in a 100-cell. Scalebar is 3 cm. Mature seeds of spring wheat (*T. aestivum* cv. Suntop) grown under LED-Supplemented glasshouse setup, UQ, Australia: **q**, plants in a 30-cell tray; **r**, plants in a 64-cell tray; **s**, plants in a 100-cell. Mature seeds of barley (*H. vulgare* cv. Commander) grown under LED-Supplemented glasshouse setup, UQ, Australia: **t**, plants in a 30-cell tray; **u**, plants in a 64-cell tray; **v**, plants in a 100-cell. Scalebar is 1 cm.

**Table 7.** Mean days to reproductive stages^3-5^ of single seed descent (SSD) sowing densities under speed breeding using the JIC-GH-LED^1^ or UQ-GH-LED^2^ protocol. JIC-GH-LED protocol used a temperature cycle regime of 22 h at 22 °C and 2 h at 17 °C to coincide with light and dark times, respectively. The UQ-GH-LED protocol used a temperature cycle regime of 12 h at 22 °C and 12 h at 17 °C.

| Species | Protocol | Sowing density | Photoperiod | Mean days to reproductive stage |
| --- | --- | --- | --- | --- |
| Spring wheat<br>*T. aestivum* | JIC-GH-LED<sup>1</sup> | 96-cell<br>(560 plants/m<sup>2</sup>) | 22 h | 45.0 ± 0.0<sup>3</sup> |
|  |  | 96-cell<br>(560 plants/m<sup>2</sup>) | 16 h | 58.0 ± 0.0<sup>3</sup> |
|  | UQ-GH-LED<sup>2</sup> | 30-cell<br>(300 plants/m<sup>2</sup>) | 22 h | 31.3 ± 0.7<sup>4</sup> |
|  |  | 64-cell<br>(640 plants/m<sup>2</sup>) | 22 h | 30.0 ± 0.0<sup>4</sup> |
|  |  | 100-cell<br>(1000 plants/m<sup>2</sup>) | 22 h | 31.0 ± 0.0<sup>4</sup> |
| Tetraploid wheat<br>*T. durum* | JIC-GH-LED | 96-cell<br>(560 plants/m<sup>2</sup>) | 22 h | 42.0 ± 0.0<sup>3</sup> |
|  |  | 96-cell<br>(560 plants/m<sup>2</sup>) | 16 h | 50.0 ± 0.0<sup>3</sup> |
| Spring barley<br>*H. vulgare* | UQ-GH-LED | 30-cell<br>(300 plants/m<sup>2</sup>) | 22 h | 27.3 ± 1.2<sup>5</sup> |
|  |  | 64-cell<br>(640 plants/m<sup>2</sup>) | 22 h | 24.7 ± 0.3<sup>5</sup> |
|  |  | 100-cell<br>(1000 plants/m<sup>2</sup>) | 22 h | 24.0 ± 0.6<sup>5</sup> |

<sup>1</sup> **JIC-GH-LED:** LED-Supplemented Glasshouse setup, JIC, UK (described in this paper, Equipment Setup Section c).
<sup>2</sup> **UQ-GH-LED:** LED-Supplemented Glasshouse setup, UQ, Australia (described in this paper, Equipment Setup Section c).
<sup>3</sup> Days to 50% ear emergence from sowing (GS55)<sup>40</sup>.
<sup>4</sup> Days to mid-anthesis (GS65) from sowing.
<sup>5</sup> Days to awn-peep (GS49) from sowing.

## 8. Acknowledgements

We wish to acknowledge the support of the Biotechnology and Biological Sciences Research Council (BBSRC) strategic programmes Designing Future Wheat (BB/P016855/1), Molecules from Nature (BB/P012523/1), Understanding and Exploiting Plant and Microbial Metabolism (BB/J004561/1), Food and Health (BB/J004545/1); and Food Innovation and Health (BB/R012512/1), and also support from the Gatsby Charitable Foundation. The benchtop cabinet was supported by an OpenPlant Fund grant from the joint Engineering and Physical Sciences Research Council/ BBSRC-funded OpenPlant Synthetic Biology Research Centre grant BB/L014130/1. SG was supported by a Monsanto Beachell-Borlaug International Scholarship and the 2Blades Foundation, AS by the BBSRC Detox Grasspea project (BB/L011719/1) and the John Innes Foundation, AW by an Australian Post-graduate Award and the Grains Research and Development Corporation (GRDC) Industry Top-up Scholarship (project code GRS11008), MMS by CONACYT-I2T2 Nuevo León (grant code 266954 / 399852), LTH by an Australian Research Council Early Career Discovery Research Award (project code DE170101296). We acknowledge Matthew Grantham and David Napier from Heliospectra for their help in the choice of LED lights, Luis Hernan and Carolina Ramírez from Newcastle University for their support and advice in the design of the benchtop cabinet, Carol Moreau from the John Innes Centre and Jaya Ghosh from the University of Bedfordshire for help with the pea and grasspea experiments, respectively, and the JIC and UQ Horticulture Services for plant husbandry and their support in scaling up SB in glasshouses.

